# Habitat heterogeneity responds to megaherbivores in East African coastal forests, but vegetation composition remains constrained by land-use history

**DOI:** 10.64898/2026.01.28.702212

**Authors:** Stella Wimmer, Elias Dauer, Jonas Eberle, Lynn Njeri, Mike Teucher, Jan Christian Habel, Maximilian Hanusch

**Affiliations:** Department of Zoology, Faculty of Science, University of South Bohemia, 370 05 České Budějovice, Czech Republic; Department of Physical Geography, Goethe-University Frankfurt, 60438 Frankfurt, Germany; Evolutionary Zoology, Department of Environment and Biodiversity, University of Salzburg, 5020 Salzburg, Austria; Wildlife Research and Training Institute, Coastal and Marine Research Center, 484-80200, Malindi, Kenya; School of Environmental and Earth Sciences, Pwani University, 195 – 80108 Kilifi, Kenya; Department of Geoecology, Institute of Geosciences and Geography, Martin Luther University Halle-Wittenberg, 06099 Halle (Saale), Germany; Institute of Landscape and Plant Ecology, University of Hohenheim, 70599 Stuttgart, Germany

**Keywords:** East African dry coastal forest, disturbance, secondary forest, Arabuko-Sokoke, tree composition, elephants, abiotic conditions

## Abstract

- Megaherbivores are increasingly promoted as agents of nature restoration, yet most research on their ecological effects has focused on temperate and non-forested systems, with limited consideration of tropical forests and their historical land-use contexts.
- A better understanding of megaherbivore impacts in tropical forests is essential to inform rewilding and restoration efforts. This is particularly important in regenerating secondary systems that historically supported megafaunga and remain highly valuable targets for ecological recovery.
- We address this knowledge gap by comparing tree species composition, forest structural attributes, and understory habitat composition across three disturbance regimes in an East African tropical dry forest: (1) primary forest with megaherbivores, (2) secondary forest with megaherbivores, and (3) primary forest without megaherbivores.
- Under megaherbivore presence, understory habitat and tree branching architecture converged across primary and secondary forests, suggesting functional consistency in disturbance effects imposed by large herbivores and indicating that key structural ecosystem processes can be rapidly restored. In contrast, canopy structure and tree species composition remained distinct between forest types and strongly constrained by persistent legacies of past human land use.
- Our findings underscore that restoration strategies relying on megaherbivores must explicitly account for historical land-use constraints rather than assuming spontaneous convergence toward primary-forest conditions.

## Introduction

Megaherbivores are ecosystem engineers that strongly affect local species inventories and habitat conditions through foraging, physical disturbance, and the redistribution of nutrients (Bakker et al., 2016; Banks et al., 2010). Consistent with these mechanisms, megaherbivore presence has been shown to increase habitat heterogeneity and promote biodiversity at both local and landscape scales (Frenette-Dussault et al., 2013; Lundgren et al., 2024; Stevens et al., 2016; Trepel et al., 2024). Consequently, the (re-)introduction of megaherbivores to degraded ecosystems is increasingly promoted as a nature-based restoration strategy across various biomes and habitats (Svenning, Buitenwerf, et al., 2024; Wang et al., 2025). However, in many ecosystems megaherbivore effects will inevitably unfold in landscapes profoundly shaped by past and ongoing human disturbance, raising the question of how far their ecological functions can operate independently of land-use legacies.

Although recent syntheses and meta-analysis show that the ecological effects of megaherbivores are broadly generalizable (Asner et al., 2009; Gordon et al., 2023; Lundgren et al., 2024), the direction and magnitude of megaherbivore impacts remain highly context dependent and often poorly predictable (Hyvarinen et al., 2021; Orr et al., 2022; Pringle et al., 2023; Trepel et al., 2024). One proposed source of this unexplained variation is anthropogenic land-use history (Schweiger et al., 2019; Trepel et al., 2024). Past and ongoing human activities restructure regional species pools, vegetation structure, and habitat conditions, thereby potentially constraining the ecological functions that megaherbivores fulfill in landscapes bearing legacies of historical land-use (Dorresteijn et al., 2015; Ellis & Ramankutty, 2008; Perino et al., 2019). Understanding how land-use history modulates the effects of megaherbivores is therefore critical for evaluating the feasibility and expected ecological outcomes of megaherbivore-based restoration measures in strongly human-modified landscapes.

Studies on megaherbivore effects have become more frequent in recent years but remain concentrated on temperate, non-forested systems and often give limited consideration to past land-use history (Monk, 2024; Svenning et al., 2016; Trepel et al., 2024). Disentangling the interactions between land-use and herbivory would be particularly critical in tropical forests, where secondary forests now account for more than half of the remaining forest area (Chazdon et al., 2009; FAO, 2020). Despite strong anthropogenic legacies, tropical secondary forests often retain substantial potential for biodiversity recovery, making them priority targets for global restoration efforts (Rozendaal et al., 2019; Strassburg et al., 2020). Most tropical forests historically supported diverse assemblages of megaherbivores, which have been extirpated across much of their former ranges (Berzaghi et al., 2018; Malhi et al., 2016; Svenning, Lemoine, et al., 2024). Therefore, a central yet insufficiently tested question is whether the effects of megaherbivores in tropical secondary forests resemble the structural and functional patterns they generate in primary forests.

The Arabuko-Sokoke Forest in coastal Kenya provides one of the rare opportunities to examine how megaherbivore impacts interact with land-use history in the tropics. The forest harbours numerous endemic and threatened species, as well as free-ranging populations of African bush elephants and African buffalo (Arabuko-Sokoke Forest Management Team, 2022). It comprises a mosaic of relatively pristine primary forest and regenerating secondary forest that originated from historical clearing for settlement in the early twentieth century and has been recovering since the late 1970s. In 2006, the installation of an electric fence excluded elephants and buffalos from parts of the forest while leaving other areas accessible, spanning both primary and secondary forest. This management intervention created a landscape-scale exclosure system encompassing three distinct disturbance regimes: (1) secondary forest with megaherbivore presence (from hereafter referred to as: *sec+*), (2) primary forest with megaherbivores (*prim+*), and (3) primary forest without megaherbivores (*prim–*). Together, this configuration enables a comparison of megaherbivore impacts across forest and historical land-use types.

Specifically, we quantified megaherbivore effects on tree species composition, canopy height and tree branching architecture, as well as understory habitat composition. Based on this data, we address the following research questions:

1. Does tree species composition differ between primary forests with and without megaherbivores, and do land-use legacies modify these patterns in secondary forests?
2. Does megaherbivore presence generate similar patterns of understory habitat composition and heterogeneity in primary forests and in secondary forests shaped by land-use legacies?
3. Does forest structure, in terms of canopy height and tree branching architecture, differ between primary forests with and without megaherbivores, and are these structural patterns modified in secondary forests by land-use legacies?

Clarifying these questions is essential for understanding how land-use history modulates megaherbivore-driven processes in tropical forests and for assessing whether these processes may contribute to the recovery of secondary forests toward less human-modified reference ecosystems (Chazdon, 2003, 2008; Galetti et al., 2017; Pringle et al., 2023)

## Methods

### Study area and plot design

Our study was conducted in the Arabuko-Sokoke Forest (ASF), a 416 km^2^ remnant of East African coastal dry forest in Kenya. The ASF has experienced a long history of human use, including settlement and intensive logging in the early 20th century, followed by protection and natural regeneration after designation as a forest reserve in 1976 (Robertson & Luke, 1993). The ASF currently supports growing populations of megaherbivores, primarily African bush elephants (*Loxodonta africana*) and African buffalo (*Syncerus caffer*). The installation of an electric fence in 2006 excluded these megaherbivores from parts of the forest while leaving adjacent areas accessible (Fig. 1), creating three distinct disturbance regimes: secondary forest with megaherbivores (sec+), primary forest with megaherbivores (prim+), and primary forest without megaherbivores (prim–).

**Figure 1.**
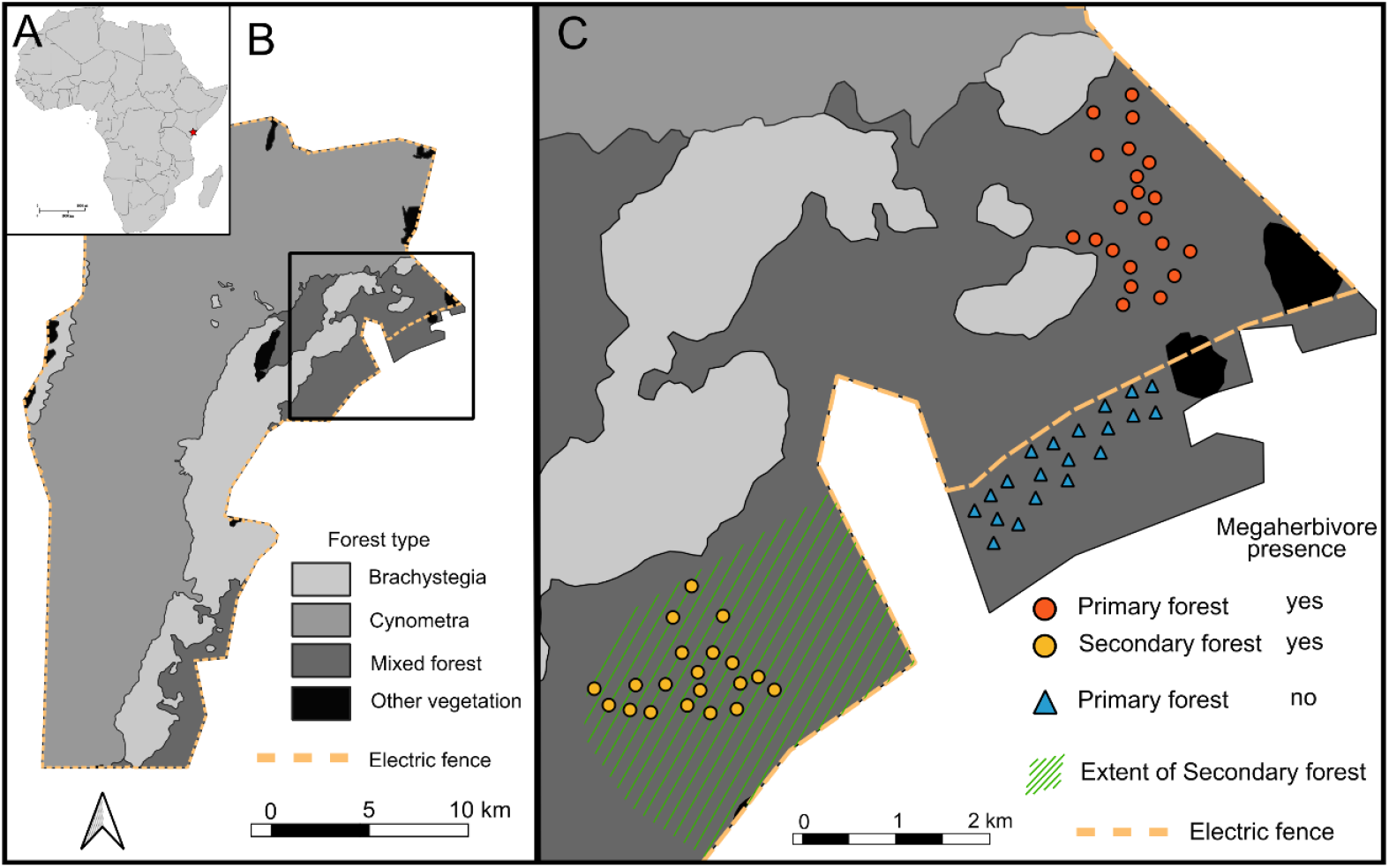
Overview of the study area. (A) Map of African continent showing the location of Arabuko-Sokoke Forest on the coast of Kenya (indicated by an asterisk). (B) Overview map of Arabuko-Sokoke Forest with major forest types delineated. (C) Overview of the study area showing the location of all sampling plots, the extent of regenerating secondary forest that was intensively logged prior to 1976, and the electric fence that creates a partial megaherbivore exclosure. All plots are situated within the same forest type (Mixed Forest).

Within these disturbance regimes, we randomly selected 60 plots (15 × 15 m each; Fig. 1), with 20 plots assigned to each regime using QGIS, maintaining a minimum distance of 100 m from roads and 200 m between plots to minimize the effects of anthropogenic influence and spatial dependence. The heterogeneous geology of coastal Kenya has given rise to multiple forest types within the area (Muriithi & Kenyon, 2002). To ensure comparability across sites, our study focused exclusively on the Mixed Forest type, which is characterised by dominant canopy species such as *Afzelia quanzensis, Hymenaea verrucosa, Combretum schumannii*, and *Manilkara sansibarensis* (Fungomeli et al., 2020; Robertson & Luke, 1993).

While this design enables a comparison across land-use histories and megaherbivore presence, we acknowledge that it does not represent a full factorial design. Despite active efforts to identify suitable sites, secondary forests without megaherbivores (i.e. *sec-*) do not exist within the study landscape. Areas outside the protected boundaries of the ASF are typically subject to intensive logging, conversion to agricultural land, or the establishment of commercial tree plantations, and no longer support either secondary or primary forest cover.

### Assessment of tree species and habitat composition

We quantified tree species composition and branching architecture by surveying all woody plant individuals with circumference at breast height (CBH) ≥ 10 cm (≈ 3.18 cm diameter) in each plot. All individuals were identified to species level, and their branching architecture was classified by the number of main stems as single-stemmed, double-stemmed, triple-stemmed, or multi-stemmed (≥ four stems). For individuals with more than one stem, CBH was measured separately for each stem; where more than four stems were present, measurements were taken for the four thickest stems (Supplementary table S1).

On each plot, we surveyed three 2 × 2 m subplots to quantify understory habitat composition and small-scale heterogeneity. Subplots were located at the plot centre and at two points positioned 12.5 m east and west of the center, respectively. Within each subplot, we visually estimated the percentage cover of herbaceous vegetation, shrubs, litter, and bare ground (Supplementary table S2). For each plot and cover component, we then calculated (i) habitat composition as the mean relative cover across the three subplots (i.e., normalized to proportions) and (ii) small-scale heterogeneity as the within-plot range (*max–min*) in relative cover across subplots.

All surveys were conducted during two sampling periods, in the dry seasons of September 2024 and February 2025, and sampling evenly split between the two periods. Fieldwork was conducted with permission from the Kenya Forest Service (permit no. KFS/GDE/R/1/VOL.1/04)

### Analysis of tree species and habitat compositon

We quantified differences in tree species and habitat composition among disturbance regimes using distance-based redundancy analysis (dbRDA; “capscale” function in the R package *vegan*, Oksanen et al., 2019), based on Bray–Curtis dissimilarities. To assess the relative contributions of megaherbivore presence and land-use history, we fitted models including both predictors and evaluated significance using 999 permutations. In addition, we tested the unique (partial) effects of each predictor by conditioning on the other predictor, allowing us to assess megaherbivore effects independent of land-use history and vice versa.

For tree species composition, we first calculated basal area for each measured stem from its CBH value. Where individuals were multi-stemmed, basal area was calculated for each stem separately and then summed to obtain an individual-level basal area. We then aggregated to the plot level by summing individual basal areas for all conspecifics within a plot, yielding plot-level basal area per species. The resulting species × plot matrix was used as the response in the dbRDA models.

For habitat composition, we used plot-level relative cover values for shrubs, herbs, litter, and bare ground (means across the three subplots) and analysed these data using dbRDA with Bray– Curtis dissimilarity, applying the same modelling and permutation framework as for tree species composition.

Differences in mean relative cover and within-plot heterogeneity among disturbance regimes were assessed separately for each cover component using non-parametric Wilcoxon rank-sum tests. Pairwise comparisons between regimes were conducted with Holm-adjusted *p*-values to account for multiple testing.

### Forest structure and branching architecture

We obtained canopy height estimates from the Meta–WRI High Resolution Canopy Height Maps (ALS–GEDI v6; registry.opendata.aws/dataforgood-fb-forests), a global 1 m-resolution canopy height product generated with machine-learning models from Maxar satellite RGB imagery and calibrated using airborne lidar canopy height maps, with additional post-processing informed by GEDI lidar observations (Dubayah et al., 2022; Lang et al., 2024). Canopy height data were accessed and processed in Google Earth Engine. We overlaid the boundaries of the plots onto the canopy height raster, extracted canopy height values for all 1 × 1 m pixels intersecting each plot. We then computed the plot-level mean canopy height that was used as a structural metric (Supplementary table S3).

Tree density (i.e. number of individual trees per plot) and branching architecture were derived from field-recorded tree data (Supplementary table S4). Branching architecture was characterised at the individual level based on the number of main stems and summarized at the plot level as the relative proportion of individuals belonging to different branching categories (single-stemmed, double-stemmed, triple-stemmed, and multi-stemmed).

Differences in canopy height, tree density, and branching composition among disturbance regimes were evaluated using non-parametric Wilcoxon rank-sum tests. Pairwise comparisons between regimes were conducted with Holm correction to account for multiple testing.

## Results

Tree species composition differed significantly among plots in relation to megaherbivore presence and land-use history, as revealed by distance-based redundancy analysis (dbRDA on Bray–Curtis dissimilarities; constrained model: *F* = 4.73, *p* = 0.001, *df* = 2, *R*^*2*^ = 0.14). Conditional permutation tests indicated that both predictors explained unique variation in species composition: land-use history (*F* = 7.19, *p* = 0.001, *R*^*2*^ = 0.11), and megaherbivore presence (*F* = 3.06, *p* = 0.004, *R*^*2*^ = 0.05). In ordination space, group separation was structured along orthogonal gradients: land-use-related differences between primary and secondary forest were primarily expressed along CAP1, whereas differences associated with megaherbivore presence were predominantly captured by CAP2.

Habitat composition also varied significantly with the imposed constraints (dbRDA on Bray– Curtis dissimilarities; *F* = 7.42, *p* = 0.001, *df* = 2, *R*^*2*^ = 0.21). However, only megaherbivore presence explained unique variation in habitat composition (*F* = 11.26, *p* = 0.001, *R*^*2*^ = 0.16), while land-use legacy had no significant independent effect (*F* = 0.56, *p* = 0.748, *R*^*2*^ = 0.02). Accordingly, ordination patterns were dominated by separation between megaherbivore-accessible and megaherbivore-excluded plots, while primary and secondary forests under megaherbivore presence showed substantial overlap (Fig. 2).

**Figure 2.**
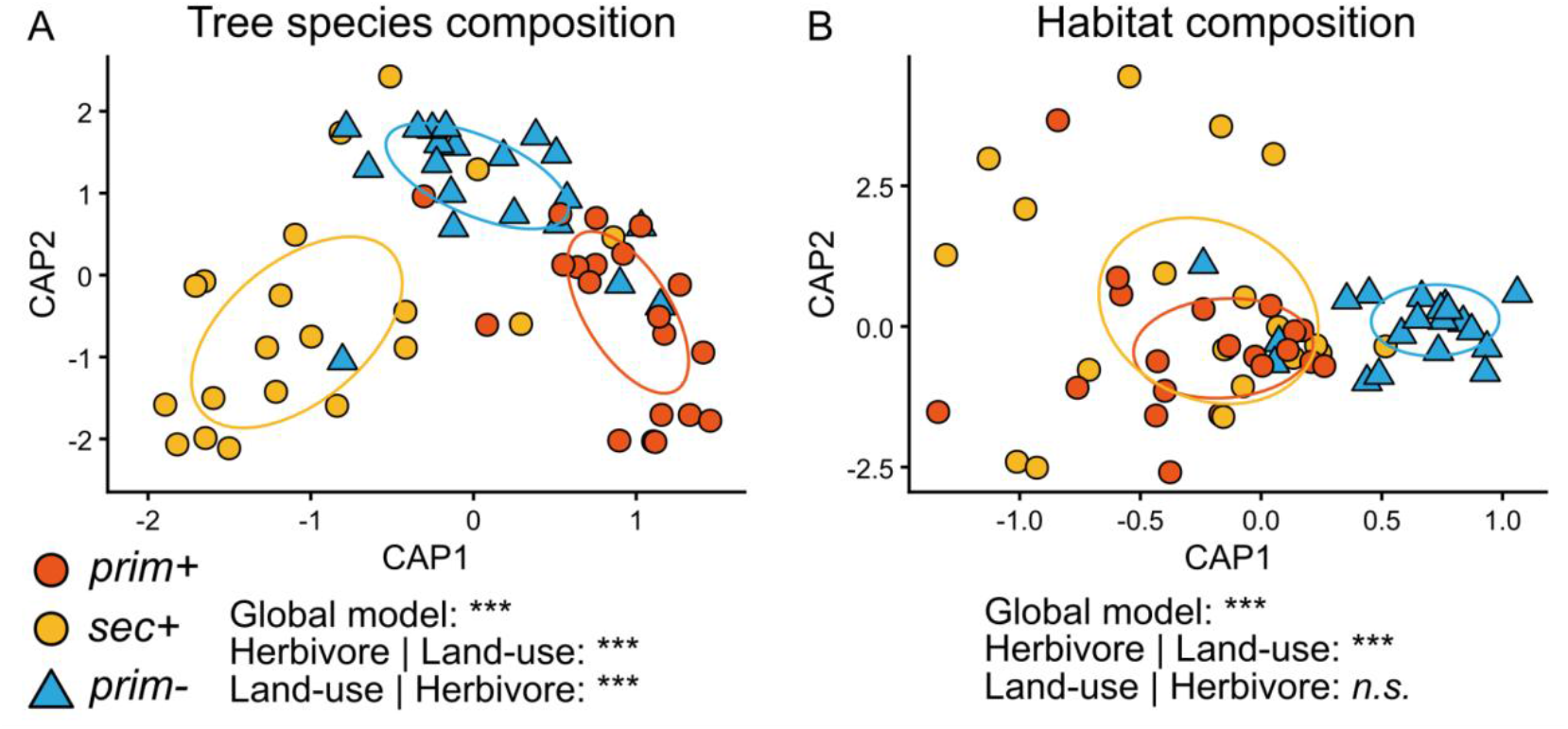
Distance-based redundancy analysis (dbRDA; Bray–Curtis dissimilarity) of (A) tree species composition and (B) habitat composition across plots. Symbols denote disturbance regimes: primary forest without megaherbivores (prim-), primary forest with megaherbivores (prim+), and secondary forest with megaherbivores (sec+); ellipses represent one standard deviation around group centroids in constrained ordination space and visualize within-group dispersion. Models were constrained by megaherbivore presence and land-use legacy; significance was assessed via permutation tests (999 permutations), and unique (partial) effects were tested using conditioned models. In (A), the global model was significant (*p* = 0.001), and both megaherbivore presence (*p* = 0.001) and land-use legacy (*p* = 0.001) explained unique variation. Differences between primary and secondary forest were expressed primarily along CAP1, whereas megaherbivore presence separated plots along CAP2. In (B), the constrained model was also significant (*p* = 0.001), but only megaherbivore presence explained unique variation (*p* = 0.001); land-use legacy had no detectable effect (*p* = 0.744). Accordingly, habitat composition was driven by megaherbivore presence, with substantial overlap between primary and secondary forest under megaherbivore access.

Individual habitat components and their small-scale heterogeneity showed pronounced variation across disturbance regimes. Shrub cover did not differ significantly among forest types (*χ*^*2*^ = 4.70, *df* = 2, *p* = 0.095; Fig. 3). In contrast, herbaceous cover differed strongly (*χ*^*2*^ = 26.3, *p* < 0.001), with significantly lower values in prim– compared to both prim+ and sec+ (*p*.*adj* < 0.001 for both), which did not differ from each other. Bare-ground cover also varied significantly (*χ*^*2*^ = 22.2, *p* < 0.001), being lowest in prim– and similarly elevated in prim+ and sec+ (*p*.*adj* < 0.001 vs. prim–). Litter cover was highest in prim– (*χ*^*2*^ = 16.1, *p* < 0.001), with lower and statistically indistinguishable values in prim+ and sec+ (*p*.*adj* > 0.9).

**Figure 3.**
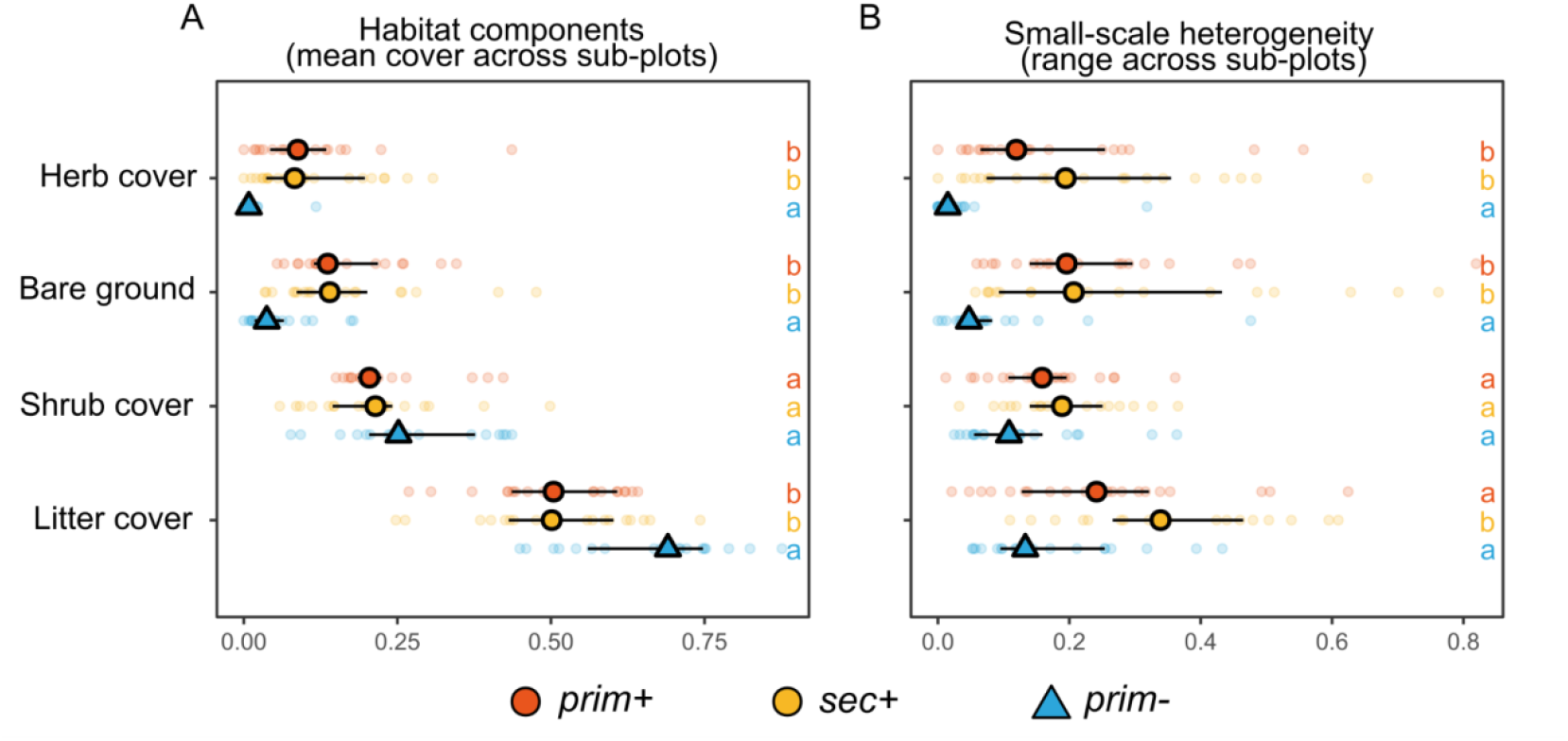
Plot-level mean relative cover of individual habitat components and their small-scale heterogeneity across disturbance regimes: primary forest without megaherbivores (prim-), primary forest with megaherbivores (prim+), and secondary forest with megaherbivores (sec+). Larger symbols indicate the median per forest type, and horizontal bars show the interquartile range. Colors denote disturbance regimes. Lowercase letters indicate significance groups from pairwise Wilcoxon tests with Holm correction; disturbance regimes sharing a letter do not differ significantly. (A) mean relative cover of shrub, herb, litter, and bare ground across three subplots per plot. (B) Within-plot heterogeneity, quantified as the range (*max–min*) of relative cover across sub-plots.

Patterns in within-plot heterogeneity mirrored these trends. Heterogeneity in herb (*χ*^*2*^ = 26.5, *p* < 0.001), bare-ground (*χ*^*2*^ = 21.7, *p* < 0.001), and litter cover (*χ*^*2*^ = 14.7, *p* < 0.001) was significantly greater in prim+ and sec+ than in prim–. For litter heterogeneity, sec+ showed the highest values (*p*.*adj* = 0.049 vs. inside). Shrub heterogeneity showed a group difference (*χ*^*2*^ = 6.42, *p* = 0.040), but no significant pairwise contrasts after correction (*p*.*adj* ≥ 0.05).

Canopy height differed significantly among disturbance regimes (*χ*^*2*^ = 31.83, *df* = 2, *p* < 0.001), with the highest values in prim+, intermediate values in prim–, and the lowest values in sec+ (all pairwise comparisons *p*.*adj* < 0.01). In contrast, tree density did not differ significantly across forest types (*χ*^*2*^ = 1.99, *p* = 0.37).

Branching architecture also varies across disturbance regimes. The relative proportion of single-stemmed individuals differed significantly (*χ*^*2*^ = 9.40, *p* = 0.009; Fig. 4), with a higher proportion in prim– compared to prim+ and sec+ (*p*.*adj* = 0.01 and 0.048, respectively). In contrast, the frequency of double-stemmed individuals showed no significant differences (*p*.*adj* > 0.55). The proportion of three-stemmed individuals differed across forest types (*χ*^*2*^ = 8.33, *p* = 0.016), with lower values in prim– than in prim+ and sec+ (*p*.*adj* < 0.05), and multi-stemmed individuals with group leaders were significantly more frequent in prim+ and sec+ than in prim– (*χ*^*2*^ = 9.41, *p* = 0.009; *p*.*adj* < 0.05).

**Figure 4.**
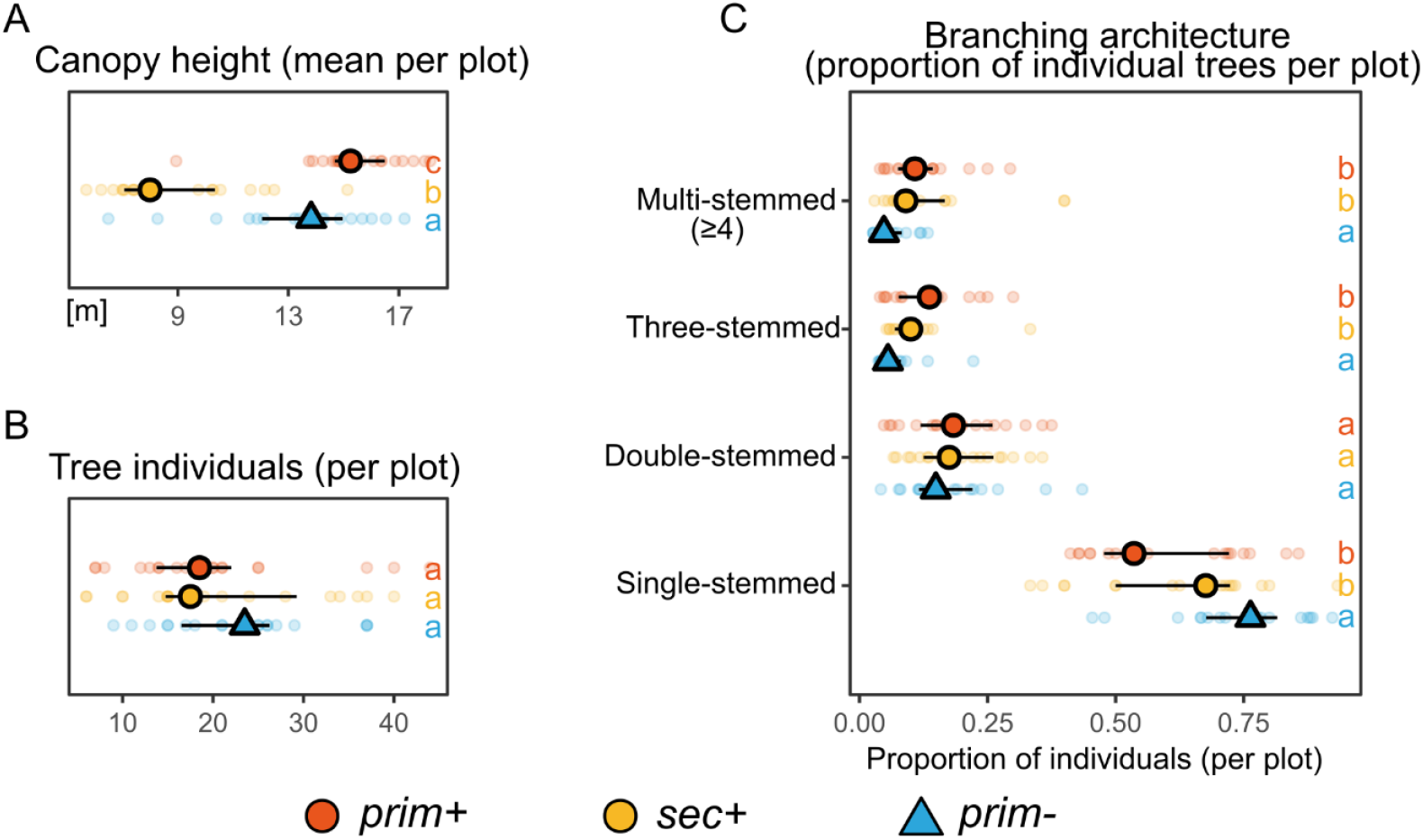
Plot-level forest structural attributes and individual tree branching architecture across disturbance regimes: primary forest without megaherbivores (prim-), primary forest with megaherbivores (prim+), and secondary forest with megaherbivores (sec+). Larger symbols indicate the median per forest type, and horizontal bars show the interquartile range. Small, transparent points represent individual plots. Colors denote disturbance regimes. Lowercase letters indicate significance groups from pairwise Wilcoxon tests with Holm correction; disturbance regimes sharing a letter do not differ significantly. (A) Mean canopy height per plot and (B) tree density (number of individuals per plot). (C) Plot-level branching architecture based on stem numbers (single, double, triple, multi).

## Discussion

Our results demonstrate that megaherbivores affect habitat composition and heterogeneity in similar ways across primary and secondary tropical dry forests. This suggests that, regardless of past human land-use, megaherbivores fulfill comparable functional roles in shaping local habitat characteristics. However, pronounced differences in tree species composition persist between disturbance regimes, even after more than five decades of secondary forest regeneration under continued megafauna presence. These enduring differences reflect long-lasting ecological legacies of historical land use that persist despite ongoing megaherbivore activity, highlighting fundamental limits to the recovery of tree species composition in tropical secondary forests.

### Lasting land-use legacies in tree species composition and stand structure

Land-use legacies exert long-lasting effects on both forest structure and species composition. Even after five decades of secondary forest regeneration in our study system (Robertson & Luke, 1993), tree species composition remains distinct from that of adjacent primary forest, and mean canopy height is consistently lower in secondary stands. This reflects incomplete structural and taxonomic recovery following past logging. Our findings align with large-scale syntheses showing that secondary tropical forests often require several decades to regain the structural complexity and species assemblages characteristic of old-growth systems (Cole et al., 2014; Rozendaal et al., 2019).

Although spatially and temporally explicit faunal baseline data for the ASF are lacking, the persistent structural differences in tree communities likely influence other trophic levels. A recent study on fruit-feeding butterflies in the same landscape revealed pronounced shifts in taxonomic and functional composition between disturbance regimes: secondary forests supported species typical of open, savanna-like habitats, while primary forests harbored taxa specialized for closed-canopy conditions (Habel et al., 2025). These findings mirror the structural differences observed in our vegetation data and highlight how incomplete structural recovery in secondary forests can cascade through trophic levels, reinforcing ecological divergence (Dent & Joseph Wright, 2009). Protecting remaining primary forest remnants is therefore critical to sustain specialist taxa that depend on old-growth habitats (DeWalt et al., 2003; Dunn, 2004).

### Effects on habitat composition and heterogeneity and branching architecture

Megaherbivores shape understory habitats in consistent ways across both primary and secondary forests in our study system. Their presence increases small-scale heterogeneity, resulting in similar habitat conditions in megaherbivore-accessible plots (prim+ and sec+). In contrast, primary forest without megaherbivores (prim–) exhibits more uniform habitat composition. This pattern suggests that megaherbivores impose a recurring disturbance regime that enhances local heterogeneity regardless of land-use history, aligning with trait-based disturbance frameworks that predict generalizable effects of large-bodied herbivores across most systems (Lundgren et al., 2024; Trepel et al., 2024). However, while habitat composition shows consistent patterns under megaherbivore presence, canopy maturity and species composition remain distinct, indicating that elephants restructure present-day habitat mosaics but cannot override the long-term ecological legacies of past land-use, even after five decades of forest recovery.

Megaherbivore presence also leads to similar tree branching architecture in primary and secondary forests. In both megaherbivore-accessible forest types, trees exhibit similar increased incidence of multiple stems, likely resulting from repeated debarking, branch breakage, and stem leader truncation (Trepel et al., 2026). In contrast, trees in megaherbivore-excluded primary forest maintain more upright, undamaged growth forms with single-stem dominance. These architectural patterns further illustrate how elephants impose a consistent structural filter across forest types, reinforcing physical habitat heterogeneity through mechanical disturbance, irrespective of successional trajectories set by past land-use.

### Constraints imposed by absence of secondary forest without megaherbivores

Our study could not include secondary forest without megaherbivores, limiting our ability to fully disentangle the effects of historical land use from those of megaherbivore presence. This constraint reflects a gap in the sampling design: secondary forest was only assessed under megaherbivore influence, and a full factorial comparison was therefore not possible. As a result, comparisons between sec+ and prim– inherently conflate land-use history with megaherbivore presence. In contrast, comparisons between prim+ and sec+ allow for a clearer assessment of land-use legacies under a shared disturbance regime. However, this limitation was not due to oversight, but to the absence of naturally regenerating secondary forests without elephants in the study landscape, where this disturbance regime no longer exists.

This limitation is not merely methodological but reflects a broader conservation asymmetry in tropical landscapes: secondary and primary forests that still support megafauna are typically located within protected areas, where forest cover and wildlife are actively maintained (ref). In contrast, forests lacking legal protection are increasingly rare, fragmented, and vulnerable to further degradation (Habel, Schultze-Gebhardt, Maghenda, et al., 2023; Habel, Schultze-Gebhardt, Shauri, et al., 2023). In southern Kenya, where our study is situated, coastal forest extent has continued to decline despite legal protection and logging bans (Teucher et al., 2020). This uneven distribution of forest types underscores how conservation infrastructure, forest persistence, and megafauna presence have become tightly coupled and now define the contexts in which forest regeneration in this landscape can be studied. However, this coupling may also form the foundation for restoring functional tropical forest ecosystems in the future.

## Conclusion

Megaherbivore research remains unevenly distributed across biomes, with a disproportionate focus on temperate and non-forested systems and limited attention to historical land-use legacies. Addressing this gap, our study shows that megaherbivore-induced disturbance creates similar habitat composition and branching architecture across forest types but does not restore the tree species inventories and some structural attributes of old-growth forest. These findings underscore the importance of integrating land-use history into restoration planning and suggest that the ecological roles of megaherbivores, while functionally consistent, are constrained by the long-term legacies of past human land-use. Yet, over successional timescales, it remains unclear whether megaherbivores facilitate convergence between secondary and primary tree species inventories. Long-term monitoring will be essential to determine whether megaherbivores promote forest recovery or reinforce a disturbance-driven feedback loop that constrains succession and stabilizes community states typical of regenerating rather than old-growth forest.

## Acknowledgements

We thank the Kenya Forest Service, Kenya Wildlife Service, and the Friends of Arabuko-Sokoke Forest for access to the forest and logistical support during field work. Furthermore, we are grateful to Mzee Geoffrey Mashauri for his invaluable help with the vegetation survey and species identification. Thanks to the anonymous referees for providing valuable feedback on a first version of this manuscript.

## Notes

### Competing Interest Statement

The authors have declared no competing interest.

